# Patterning on the move: the effects of Hh morphogen source movement on signaling dynamics

**DOI:** 10.1101/2021.06.04.447051

**Authors:** D. G. Míguez, A. Iannini, D. García-Morales, F. Casares

## Abstract

Morphogens of the Hh-family trigger gene expression changes of receiving cells in a concentration-dependent manner. The outputs of the pathway include regulation of cell identity, proliferation, death or metabolism, depending on the tissue or organ. This variety of responses relies on a conserved signaling pathway. Its internal logic includes a negative feedback loop involving the Hh receptor Ptc. In this paper, we use experiments and computational models to study and compare the different spatial signaling profiles downstream of Hh in several developing *Drosophila* organs. We show that the spatial distribution of Ptc and the activator form of the Gli transcription factor, CiA, in wing, antenna and ocellus show similar features, but markedly different from that in the compound eye (CE). We show that these two profile types represent two time points along the signaling dynamics, and that the interplay between the spatial displacement of the Hh source in the CE and the negative feedback loop maintains the receiving cells effectively in an earlier stage of signaling. These results indicate that the dynamics of the Hh source strongly influences the signaling profile Hh elicits in receiving cells, and show how the interaction between spatial and temporal dynamics of signaling and differentiation processes can contribute to the informational versatility of the conserved Hh signaling pathway.

## INTRODUCTION

The development of almost all organs of any animal relies on Hh signaling for the control of their growth, differentiation and/or patterning. These roles extend beyond early development into the adult, where the Hh pathway is required for tissue homeostasis and regeneration. In addition, mutations affecting components of this pathway contribute to a number of congenital diseases and cancer types (Ingham et al., 2011). Although the biochemical and cellular details of Hh ligand production, distribution and receptor-binding, and signal transduction events downstream of Hh are extremely intricate (and to date not fully elucidated), the Hh signaling pathway seems to be governed by a universal underlaying regulatory logic ((Kong et al., 2019; Li et al., 2018) and Figure 1). The major output downstream of Hh is the regulation of gene transcription in receiving cells (Briscoe and Therond, 2013; Ingham et al., 2011; Lee et al., 2016; Petrova and Joyner, 2014). The transcription factors in charge of this transcriptional regulation belong to the Gli family, which in *Drosophila* are represented by the single *ci* (*cubitus interruptus*) gene. In the absence of signal, Ci/Gli undergoes proteolytic cleavage into a transcriptional repressor (in *Drosophila* it is named “CiR”). Hh, binding to its receptor Ptc/PTCH, relieves Ptc repression on Smo. Free from this repression, Smo blocks the proteolysis of Ci/Gli to render a full length Ci/Gli that acts as a transcriptional activator (“CiA” in *Drosophila*) (Aza-Blanc et al., 1997). Interestingly, Ptc is among the transcriptional targets of the pathway: cells receiving Hh up-regulate *ptc* expression (Figure 1; (Capdevila et al., 1994; Tabata and Kornberg, 1994). As a result of this feedback, increasing levels of Ptc, by binding to Hh, dynamically alter the quantitative profile of Hh across the field of Hh-receiving cells and modify the signaling levels in those cells (Briscoe et al., 2001; Garcia-Morales et al., 2019; Li et al., 2018; Miguez et al., 2020). Hh proteins are produced in specific domains within a developing organ and lay adjacent to Hh-responsive cells expressing *ptc* and *ci*. This Hh-producing cells are often insensitive to it (for example, in many *Drosophila* organs, Hh-producing cells express the transcriptional repressor *engrailed (en)* which turns-off the expression of *ptc* and *ci* (Sanicola et al., 1995; Schwartz et al., 1995)). Responsive cells alter their gene expression according to the intracellular signaling levels elicited by the concentration of Hh they receive. As a result, different gene expression patterns are elicited at different distances from the Hh source, resulting in patterned cell behavior. These are the characteristics defining a “morphogen” (as defined by L. Wolpert in (Wolpert, 1969)). This morphogen-type action has been characterized in a number of systems, including the *Drosophila* wing primordium-usually called wing “disc” (Mullor et al., 1997; Strigini and Cohen, 1997).

**Figure 1:**
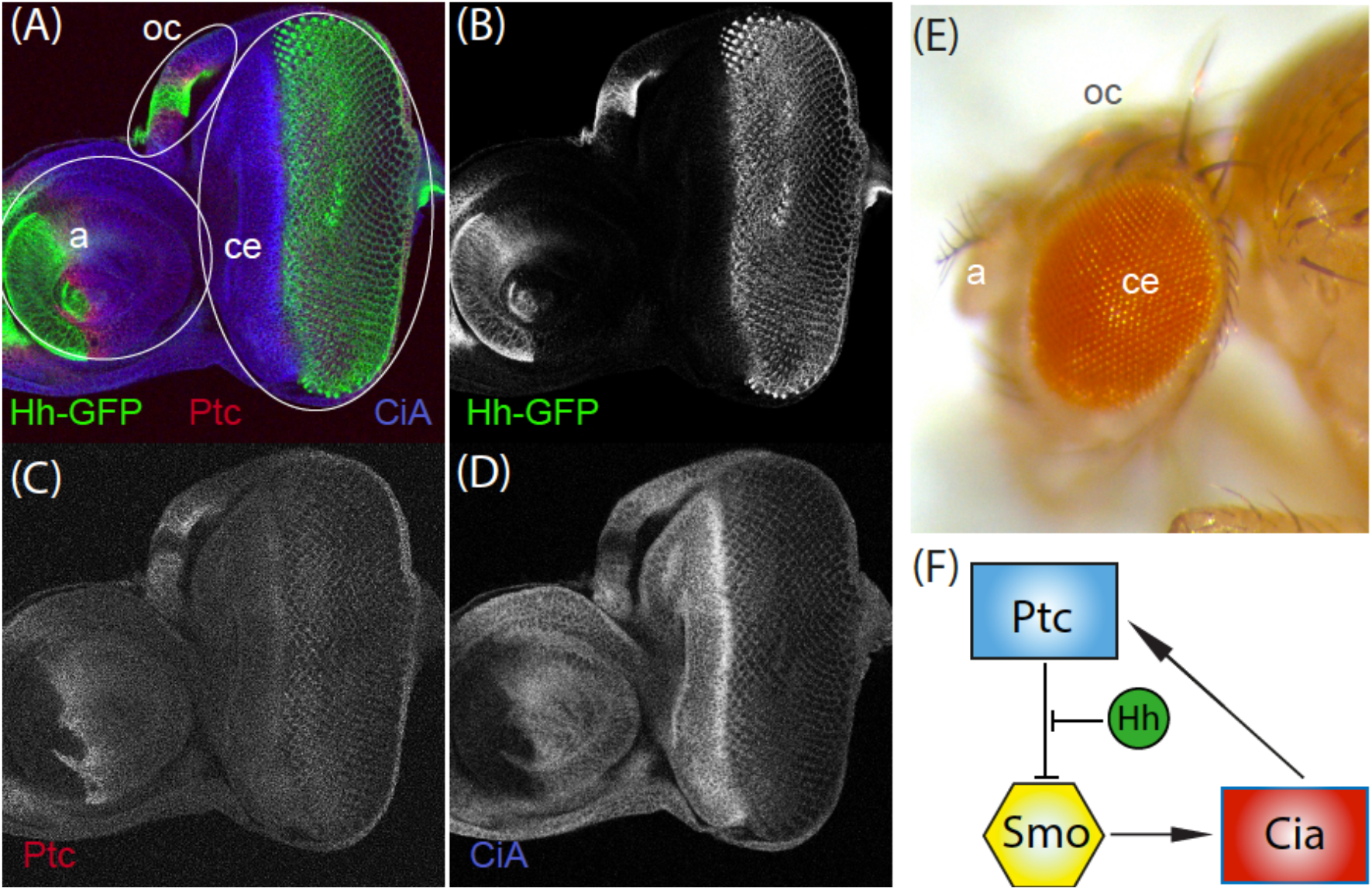
(A-D) Confocal image of a Hh:GFP eye-antennal disc stained for GFP, Ptc and CiA. Merged (A) and individual signals (B-D) are shown. (E) Adult *Drosophila* head showing the adult derivatives of the eye-antennal disc: compound eye (“ce”), antenna (“a”) and ocelli (“oc”). (F) Basic architecture of the Hh signaling pathway. In the absence of Hh signal, Ptc inhibits the production of CiA by Smo. By binding to Ptc, Hh relieves this inhibition. The production of CiA results in the transcriptional activation of the pathway’s targets. One of these targets is Ptc itself.

Another organ in *Drosophila* that requires Hh for its development is the *Drosophila* compound eye (CE). In this organ, once the first photoreceptor R-cells differentiate, they start expressing Hh. Hh, dispersing from these R-cells, activates the expression of the proneural gene *atonal (ato)* in cells immediately anterior to them. *ato* then triggers a series of regulatory steps that lead ultimately to the differentiation of new R-cells (Dominguez, 1999; Jarman et al., 1995). As part of their differentiation process, these new R cells become sources of Hh as well. This positive feedback loop results in a wave of differentiation that sweeps across the eye primordium until the pool of progenitor cells is exhausted (REVIEWED in (Miguez et al., 2020)), in the form of a moving wave-front characteristic of positive feedback loops (Miguez et al., 2007). Therefore, in its eye function, rather than generating spatially patterned cell diversity, Hh reiteratively induces only one major cell fate transition (a proneural fate) along developmental time. This feature, together with the fact that the source of Hh is moving as differentiation proceeds, contrasts with the situation in the wing, where the source of Hh is spatially static and acts as a prototypical morphogen.

This inherent versatility of the Hh pathway, that allows it to drive the differentiation of organs as different as wings, eyes or antennae is still not fully understood. In this manuscript, we combine modeling and experiments to investigate this question by comparing the dynamics and signaling profiles downstream Hh in different organs of the developing *Drosophila*. Our results suggest that the interplay between the dynamics of the negative feedback between Hh and Ptc and the dynamics of the Hh source provide the versatility to produce the different signaling signatures required to drive patterning in different organs.

## RESULTS & DISCUSSION

We start our analysis with a quantitative characterization of the Hh signaling in different developing organs in the fly. To do this, we used confocal microscopy to measure simultaneously the levels of Hh (signal) across the fixed eye-antennal and wing discs of late third-stage larvae, together with the expression profiles of two signaling readouts: Ptc (receptor, pathway repressor and pathway transcriptional target) and the activator form of CiA (transcriptional activator) (Figure 1). The eye-antennal disc gives rise to the CE as well as the antenna and the ocelli (small single-lens eyes located on the head’s forefront) (Figure 1; (Haynie and Bryant, 1986)). Each of these organs expresses Hh (Figure 1) and requires Hh signaling for their development (Heberlein et al., 1993; Ma et al., 1993; Mohler, 1988; Royet and Finkelstein, 1997). To obtain a functional Hh profile, the larvae carried a BAC containing a functional GFP-tagged Hh gene (Chen et al., 2017), so that the GFP signal is a faithful reporter of Hh protein distribution. Ptc and the transcriptional activator form of Ci (CiA) were detected using specific antibodies (See Materials and Methods) (Figure 2). In all four organs, the Hh:GFP signal describes a declining profile from the source, as expected. However, we noted two things. First, the signaling profiles of Ptc and CiA in the wing, antenna and ocellus were qualitatively similar, with a first peak of Ptc followed by a peak in CiA further from the Hh source. On the other hand, the signaling signature of Hh in the CE was qualitatively different from the others, with a peak expression of Ptc that is coincident in space with a maximum in CiA. A second main difference is that the signal intensity of Ptc at this maximum was significantly lower in the CE that in the other tissues.

**Figure 2:**
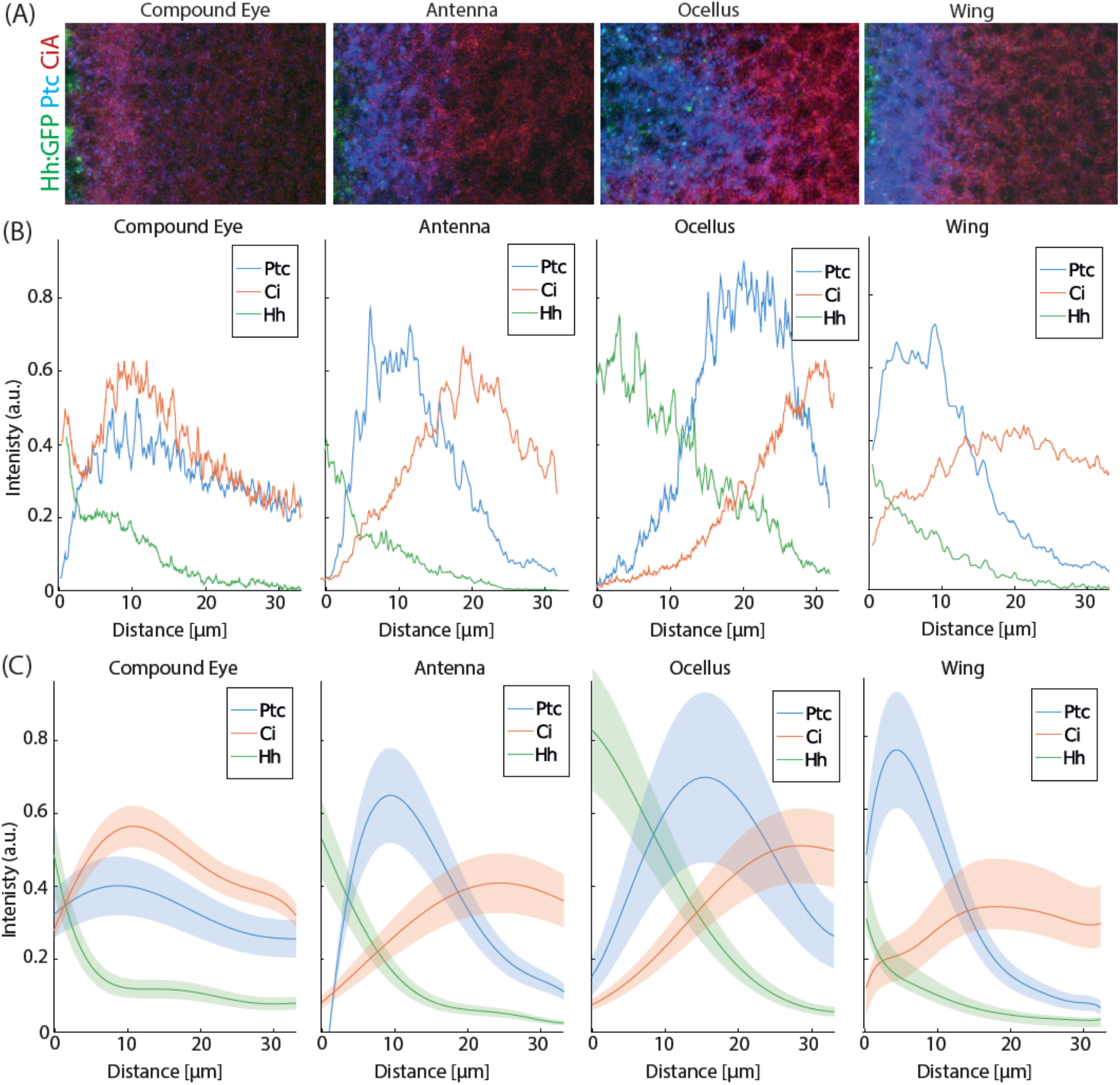
(A) Representative snapshots of Compound eye, Antenna, Ocellus and Wing disc stained for Ptc (blue), CiA (red) and Hh:GFP (green), spanning the region abutting the Hh-expressing domain. (B) Representative values of the spatial distribution of the levels of Ptc (blue), CiA (red) and Hh (green) for the four snapshots above. The Hh:GFP source is located at distance = 0. (C) Average values of the different staining profiles for at least five independent replicates for each tissue (EXTENDED DATA Figure 2: spreadsheet table with measurements used). Ribbon indicates the standard error of the mean value.

The observation of non-coincident peaks for CiA and Ptc in wing, antenna and ocellus was counterintuitive at first: With increasing Hh concentrations received, the levels of CiA produced should also increase. Since Ptc is a transcriptional target of CiA, both their profiles should follow each other -i.e. their maxima should coincide, as observed in the CE. The observation of lower levels of Ptc relative to CiA in the CE were not easily explained either. For example, high Ptc relative to CiA was observed in the antenna as well as in the ocellus, where maximum Hh levels at the source were very different, while CE and wing, with similar Hh levels at the source, show low and high Ptc/CiA signal ratios, respectively. One possibility to explain these organ-specific differences in signaling profiles was that the Hh-signaling core pathway is subject to organ-specific modifications. An alternative explanation is that the two signaling signatures of Hh can emerge without the need of organ-specific modifications. To explore this second possibility, we built a computational model of the Hh core signaling pathway, as outlined in Figure 1, and modeled as indicated in Figure 3.

**Figure 3:**
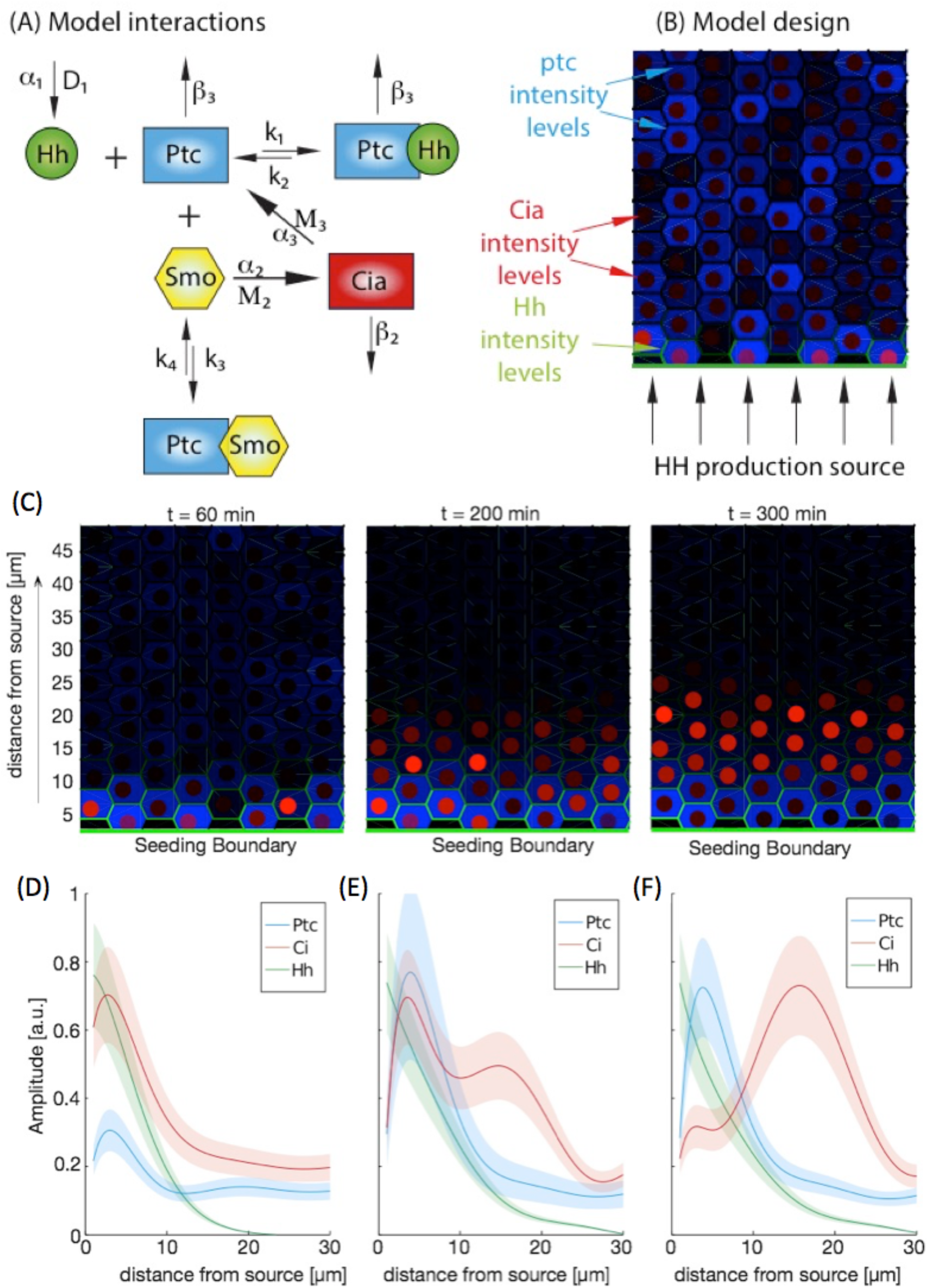
(A) Scheme of the interactions of the numerical model, with the parameters used. (B) Spatial configuration of the model as a two-dimensional array of 10×10 cells. the Hh source is located at the lower boundary. Protein levels in each cell are expressed in colors. For visualization purposes, we represent Ptc levels in blue in the cytoplasm, Hh levels in green at the membrane, and CiA levels in red in the nucleus. (Extended DATA Figure 3: code of 2D model). (C) Output of a single simulation at different times, in terms of the total levels of Ptc, CiA and Hh at different time points. (D-E) spatial profiles of Ptc, CiA and Hh computed based on distance of each cell to the Hh source. Lines represent average values of fitted curves for 5 independent numerical simulations. Ribbons represent 50% confidence interval.

The model is first formulated as a system of ordinary differential equations (ODEs) that capture the core pathway: The binging of a diffusible Hh to its receptor Ptc relieves the repression that Ptc exerts on Smo. This latter, when freed, activates Ci to produce the transcriptional activator, CiA which, in turn, feeds back onto the pathway by activating Ptc production (Figure 3A). The systems of equations is:

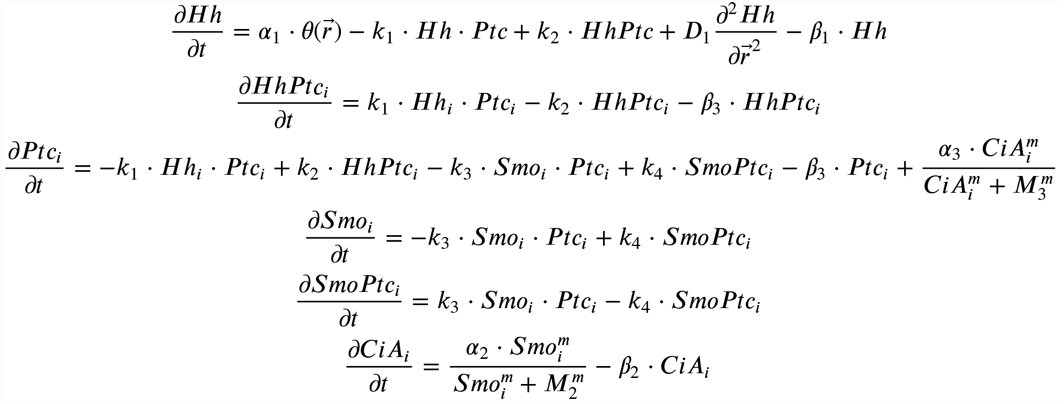

The first equation represents the binding and unbinding of free Hh and free Ptc (rate constants k_1_ and k_2_) forming the complex HhPtc. The ø(t) function indicates that Hh is produce at rate α_1_ only at specific locations in the system (the lower boundary). D_1_ is the diffusion coefficient and ß_1_ is the constant degradation of the free molecule.

The second equation models the formation dynamics of the complex between Hh and Ptc, with degradation constant ß_3_. The third equation corresponds to the dynamics of free Ptc, which can also bind and unbind to Smo (rate constants k_3_ and k_4_), and degradation constant ß_3_ (similar to free Ptc). The last term represents the activation of free Ptc production by CiA as an activating Hill function with Hill coefficient m and constant M_3_. The fourth and fifth equations correspond to the dynamics of free (“Smo”) and bound (“SmoPtc”) Smo. The final equation illustrates the production of CiA by Smo, also as a simple Hill function with rate constant α_2_ and Hill coefficient m and constant M_2_. CiA is assumed to degrade with rate constant ß_2_.

To study the generation of the signaling profiles in the tissue, we have extended our model as a spatial and temporal hybrid by embedding the ODEs in a spatial array of 10X10 hexagons representing the bidimensional epithelial cell layer of the fly organs. An example of this computation is shown in Figure 3B. To reproduce the biological situation, Hh enters the system from the bottom boundary and then diffuses across the tissue. The temporal scale was calibrated using measurements by Nahmad and Stathopoulos (Nahmad and Stathopoulos, 2009), in which the authors meaured the time required for maximal activation of Ptc by Hh in the wing disc (360 min at 18°C). Since developmental time at 18°C approximately halves at 25°C (temperature at which our experiments were carried out)(Al-Saffar et al., 1995), the model parameters were calibrated to produce maximal Ptc activation at around 180 min after Hh stimulation (see Materials and Methods).

When we ran the simulations and follow the profiles over time (Figure 3C, and Supplementary Movie 1), we observe that at earlier times the Hh gradient that is formed drives CiA and Ptc with peaks in similar positions, close to the gradient’s source. However, as time proceeds, CiA maximum is displaced as Ptc expression increases close to the source (Figure 3D-F). Therefore, the model reproduces the profiles for the CE and the other organs as an early and a later time point, respectively, along its temporal dynamics. The peak displacement can be explained by our model as a consequence of the delay between CiA activation and the feedback mediated by Ptc: initially, Hh reception leads to CiA production followed by an increase in Ptc transcription. But, as time passes and Ptc expression increases due to CiA, its negative action on the pathway (inhibiting Smo) results in the reduction of CiA levels close to the source where Ptc increases. As the Ptc profile sharpens close to the source, the non-bound Hh ligand keeps dispersing farther and activates CiA, which now shows a peak beyond that of Ptc. At this point the system reaches a quasi-stationary state that strongly resembles the experimental condition measured in the wing, antenna and ocellus (Figure 2C).

These results pointed us to further investigate why the CE’s signaling profile appears to be at an earlier stage of this dynamics. As discussed in the previous sections, the Hh source is spatially static in wing, antenna and ocellus, while it moves in the CE. If the receiving cells, as a result of the differentiation wave, became a Hh source faster than the time required for the activation of the negative feedback loop that separates the Ptc and CiA peaks, the Hh signaling profile would always resemble an “early stage” signaling profile. This would imply that different signaling profiles in different organs could simply emerge from the signaling dynamics and no organ-specific modifications of the pathway would be needed to explain them. To test this hypothesis, we carried out an experiment in which we delayed the movement of the differentiation wave. To do it, we expressed an RNAi targeting specifically *ato* in the developing CE for a period of 20 hours, after which we dissected the eye-antennal discs and profiled the expression of CiA and Ptc (See Materials and Methods). By attenuating *ato*, the process of R cell recruitment should be delayed or halted, as *ato* is required for the specification of the founder R cells, the R8 type (Jarman et al., 1995), which are responsible of the induction of all other retinal cells, including the remaining R cell types which express Hh (reviewed in (Roignant and Treisman, 2009). Indeed, the adults from this experiment showed reduced eyes (Figure 4). When we compared the CiA and Ptc profiles in control and *ato-*RNAi (“mutant”) discs, the profiles of mutant CEs were intermediate between those of control CEs and of antennae (as the antennae were not affected by the eye-specific manipulation, they showed profiles indistinguishable among control and mutants)(Figure 4A): In the *ato-*RNAi CE the Ptc profile showed increased maximal amplitude and a steeper slope, both differences statistically significant, plus a slight displacement of the CiA peak compared with control CEs (Figure 4B,C). This profile is equivalent to the computational profile at intermediate times and therefore supports our hypothesis that the two types of signaling profiles we have observed arise from the same signaling network and depend on whether the Hh source is spatially static or not.

**Figure 4:**
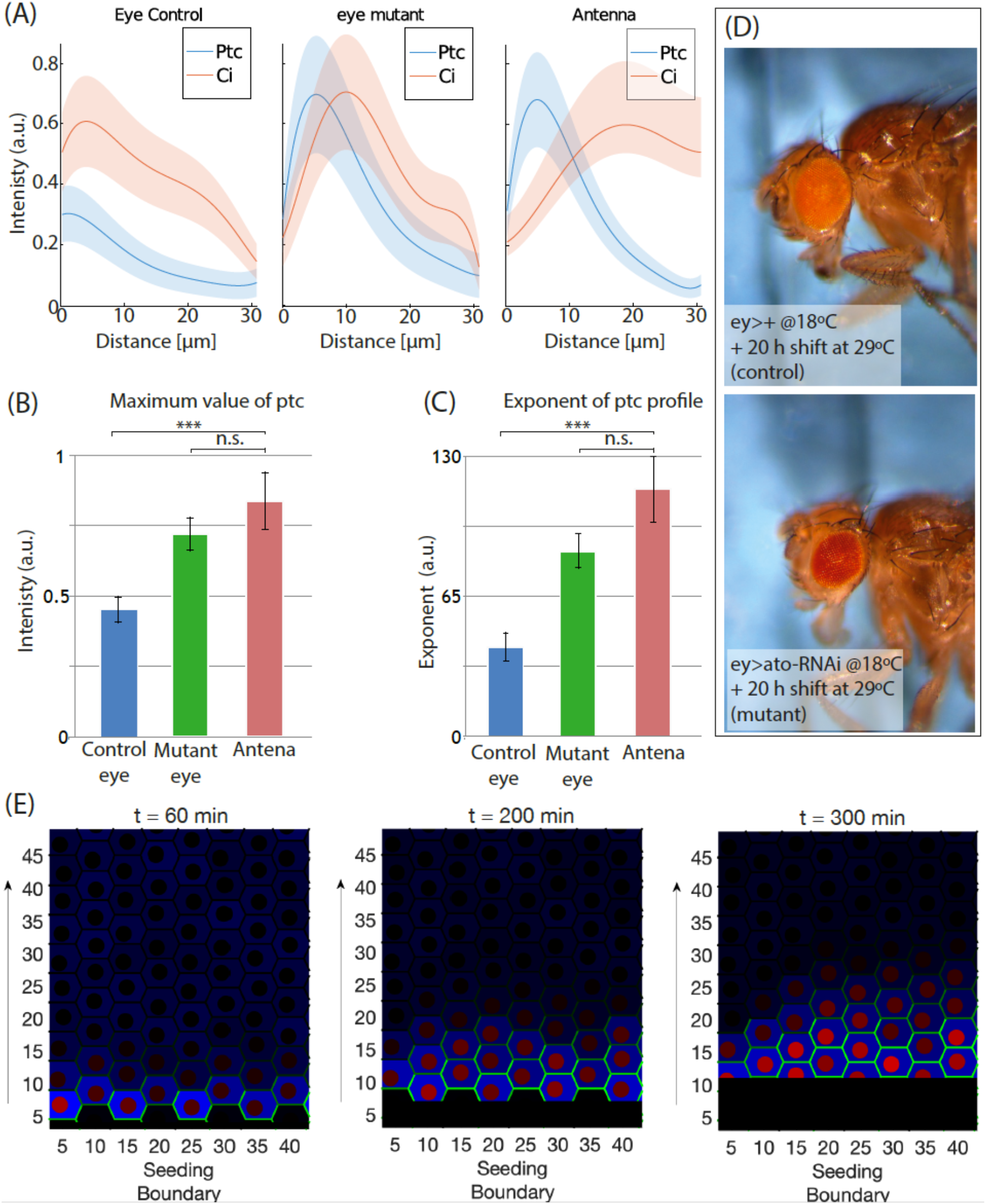
(A) Average spatial distribution of the different profiles of Ptc and CiA (at least four independent replicates) for control and mutant compound eye and antenna. (B) Quantification of maximum Ptc values in these three conditions. (C) Quantification of exponent of the exponential fitting of the decaying region in the Ptc profile, for the three conditions tested. Statistical analysis (Student’s test, two sample, unequal, one tail) shows that the differences between the mutant and the antenna are not-significant, but highly significant when the control and mutant CEs are compared. Error bars indicate the standard error of the mean value. (D) Adult control (above) and *ato-*mutant (below) flies. The transient stalling of the differentiation process results in smaller eyes. See main text for detailed explanations of the experiment. (E) Computational experiment in which the Hh producing front moves. As in Figure 3, Hh is green, Ptc is blue and CiA is red. Snapshots at 60, 200 and 300 min. The maxima of CiA and Ptc expression are coincident even at 300 min.

To test this hypothesis further, we used again our computational model. In this experiment, we repeat the same conditions and parameters values, and the only difference is that now we set the Hh source to move across the cell field at a speed of one cell per hour (the reported velocity of the signaling wave, at 25°C is about 3,4μm/hour (Vollmer et al., 2016; Wartlick et al., 2014), with 3-4μm being the approximate diameter of an epithelial disc cell). Figure 4B,C shows three snapshots of this scenario taken at three different time points. A time lapse movie is presented as Supplementary Movie 2, to be compared with the scenario of static Hh source presented in Supplementary Movie 1). The simulation shows that, when the source of Hh is moving across the tissue, the separation between the peaks of CiA and Ptc does not occur, which recapitulates the profiles obtained experimentally in the CE (Figure 2A). This contrasts with the result of a static source, where the same parameters and conditions result in a clear separation of the maxima at 300 min (Figure 3C-D, right panels), similarly to the profiles of antenna, ocelli and wing primordia (Figure 2C).

In conclusion, we show that these different Hh signaling profiles are just different states along the pathway’s temporal dynamics. In the CE, Hh-receiving cells start to differentiate into Hh-producing cells faster than the time required for downstream interactions to reach a steady state, which is attained in other tissues in which the Hh source is spatially static.

The two signaling profiles that we have described, one characterizing the CE and the other shared by all the other organs analyzed, likely differ in the information they carry. The wing/antenna/ocellus profile defines two clearly different signaling peaks which may be further enriched by the temporal dynamics of the signaling before reaching a steady state. In the CE, though, the profiles suggest that once cells start receiving Hh signaling, they increase their signaling until they are taken over by the differentiation wave (and initiate their differentiation that transform them from Hh-receiving to Hh-producing). Therefore, the difference in signaling profiles we have noted may thus reflect different functional “modes”. In the “spatial mode” Hh generates an information-rich, spatially static gradient that allows the definition of several distinct positional values, such as those Hh instructs in the wing and therefore works as a morphogen. In the CE, Hh works in a “mobile mode”: The moving gradient is less rich in spatial information as it is used to trigger a single cell fate repeatedly across a field of cells. In this mode, Hh works as a fate “inducer”, rather than as a prototypical morphogen. In this mobile mode, the speed at which the Hh source travels depends on the kinetics of the biochemical steps triggered by the reception of the signal initiating photoreceptor cell differentiation and their production of Hh, resulting in the coupling between gradient movement and cell differentiation.

## MATERIALS AND METHODS

### *Drosophila* strains and genetic manipulations

The Hh:GFP strain is described in (Chen et al., 2017). They carry a BAC chromosome harboring a functional tagged *hh* locus. This is a rescuing line -i.e. it is able to rescue a null *hh* genetic condition and therefore Hh:GFP can be confidently considered a good reporter of the normal Hh protein distribution.

To attenuate *atonal (ato)*, larvae from the cross *ey-GAL4* (Flybase: https://flybase.org/reports/FBtp0002646) females to *UAS-atonal RNAi* (VDRC 48675; (Dietzl et al., 2007)) males were raised at 18°C for most of their development. The GAL4/UAS system is temperature sensitive, with increasing expression at higher temperatures. At 18°C, the eyes of the resulting adults were indistinguishable from controls (not shown). To induce the expression of the *ato*-RNAi construct, larvae were transferred at 29°C and dissected 20h afterwards. We also performed the analysis at later times (~30h at 29°C), but at this point the CiA and Ptc signals were very low anterior to the stalled retina, likely because of increased cell death (not shown), so the profiles were not used. Adult flies were imaged using a Leica 490 digital camera on a Leica DFC 320 binocular scope.

### Confocal microscopy and image analysis

Immunolocalization and confocal microscopy was done as described in (Garcia-Morales et al., 2019). Antibodies used were: rabbit anti-GFP (A11122, at 1/1000; Molecular Probes); mouse anti-Ptc (Apa1; at 1/100) and rat anti-CiA (2A1; at 1/5) were obtained from the Developmental Studies Hybridoma Bank, created by the NICHD of the NIH and maintained at The University of Iowa, Department of Biology, Iowa City, USA. Alexa-fluor-conjugated secondary antibodies, diluted at 1/400, were from Molecular Probes. Confocal images were collected as z-stacks and processed using Fiji (Schindelin et al., 2012). Strips of tissue were oriented such that they were orthogonal to the border of the Hh source, which in the CE corresponds to the morphogenetic furrow (MF). These strips were further trimmed to exclude source cells. Then, 2 to 4 optical z-sections, selected as containing most of the signal per oriented and trimmed stack were projected using the “maximal projection” tool and the signal for each of the three channels (each representing one of the three proteins, Hh:GFP, Ptc and CiA) was measured and extracted as a spatial profile.

### Computational model

Numerical simulations of the model equations were performed using a computational script written in Matlab© (The Mathworks) and developed in house (available as source code File 1). In the model, space and time are discretised using the Euler algorithm. Concentrations are considered as dimensionless, but space and time are dimensional (μm and minutes, respectively). The model consists on a hybrid approach that combines ordinary differential equations (ODEs) for the processes inside cells and partial differential equations (PDEs) for processes across cells, such as morphogen transport. Cells are simulated as two-dimensional hexagonal regions in a Voronoi diagram.

The simulation process is as follows: values for the initial concentrations of the variables and for each of the model parameters and for each cell “i” in the tissue are obtained from a gamma distribution with standard deviation set to 10% of the mean value, to represent cell variability (mean value for each parameter is listed in table 1). For each time step, the first equation for Hh diffusion is integrated globally in space and time. Due to the spatial gradient, the amount of Hh that each cell is receiving, Hh_i_, depends on its position. This value Hh_i_ is computed for each cell and time step, as the average concentration for all pixels inside each cell. Next, the value of Hh_i_ is used to integrate the six ODEs (see equations in the main text) independently for each of the 100 cells in the system. Finally, to obtain the profiles of expression levels, the values for the total amount of each variable are computed and averaged based in the distance of each cell “i” to the source (located at x=0). This sequence of events is performed for each time point during the simulation.

The source of Hh is set in our system as the lower boundary (labelled as Seeding Boundary in the plots). From here, Hh enters the spatial system and its transport is simplified as standard two-dimensional diffusion. In numerical simulations that involve a moving morphogenetic furrow, this is introduced by translocation of the lower boundary of the system (that is continuously secreting the Hh protein). This way, the boundary is set to move at constant speed upwards traveling across the system. Cells that have been “run over” by the moving furrow boundary are set to differentiate and stop consuming Hh (colored in black in the simulation, for visualization purposes).

### Spatial and temporal calibration

To reduce the number of free parameters and equations, and to obtain a simpler model that could be more easily explored, we opted to condense several features of the Hh pathway in a small set of interactions. Therefore, some parameter values cannot be directly informed from the literature, because they encompass multiple interactions. The correct ranges of some parameter values have to be estimated by comparison with the dynamics and steady state values of the real system.

Table 1 shows the values of the model parameters used in our simulations. For the spatial calibration, we estimated the size of a cell at around 5 μm in diameter (estimated from the experimental snapshots).

For the Hh diffusion, we use a value D of 0.1 μm^2^ /min similar to the values obtained from experimental measurements in the literature, and that produces an exponential profile of characteristic length of around 12 μm in steady state. We have measured previously that the characteristic length of the Hh exponential profile is around 13 μm and that is conserved for antenna, ocellus, compound eye and wing disk (Miguez et al., 2020). This value of Hh diffusion, combined with the value of Hh production at the source and degradation rates also result in a steady state configuration reached at around 60 min in the absence of other downstream interactions, which is consistent with our previous data (Garcia-Morales et al., 2019).

The calibration of the time variable in the model is based on experimental data from (Nahmad and Stathopoulos, 2009). Here, authors estimate that Ptc levels regain their maximum after 360 min at 18°C. Dynamics at standard culture conditions (25°C) is twice as fast as the dynamics at 18°C, so we estimate that the increase of Ptc in standard conditions occurs at 180 minutes. Based on this data, we set our parameter values to obtain a peak in Ptc levels at around 180-200 min after Hh stimulation. This time reference is used to estimate the equivalence between time step in the simulation and minutes in the experimental system.

Finally, affinity and dissociation of Hh and Ptc has been estimated from previous experimental data, were we estimated that, in steady state conditions, around 90% of the Ptc is bound to Hh, and that this percentage is independent of the distance to the Hh source (Miguez et al., 2020).

| Parameter | explanation | Value | units | source |
| --- | --- | --- | --- | --- |
| $D_1$ | Diffusion constant of Hh | 0.1 | $\mu\text{m}^2/\text{min}$ | (Garcia-Morales et al., 2019) |
| Time | Length of simulation | 300 | min |  |
| Cell diameter | Size of a progenitor cell in the tissue | 5 | $\mu\text{m}$ | Exp data |
| Furrow velocity | Speed of the moving boundary in the CE | 5 | $\mu\text{m}/\text{min}$ | (Vollmer et al., 2016) |
| $k_1$ | Rate of Hh and Ptc binding | 0.15 | $1/(\text{Conc} \cdot \text{min})$ | Estimated from exp data |
| $k_2$ | Rate of Hh and Ptc unbinding | 0.01 | $1/\text{min}$ | Estimated from exp data |
| $\alpha_1$ | Level of Hh at source | 300 | conc | calibrated |
| $\beta_1, \beta_3$ | Degradation of Hh, Ptc and HhPtc complex | 0.001 | $1/\text{min}$ | calibrated |
| $k_3$ | Rate of Ptc and Smo binding | 0.03 | $1/(\text{Conc} \cdot \text{min})$ | (Nahmad and Stathopoulos, 2009) |
| $k_4$ | Rate of Ptc and Smo unbinding | 0.01 | 1 / min | (Nahmad and Stathopoulos, 2009) |
| $m$ | Hill exponent | 5 | nondimensional | calibrated |
| $\alpha_2$ | Production of Ci by Smo | 2 | Conc / min | (Nahmad and Stathopoulos, 2009) |
| $\beta_2$ | Degradation rate of Ci | 0.04 | 1 / min | (Nahmad and Stathopoulos, 2009) |
| $\alpha_3$ | Production Ptc by Cia | 0.2 | Concentration/min | (Nahmad and Stathopoulos, 2009) |
| $M_2$ | Hill Coefficient of Ci expression | 2.5 | Concentration | estimated |
| $M_3$ | Hill Coefficient of Ptc expression | 5 | Concentration | estimated |

### Statistical Analysis

Average profiles for the expression levels of Hh, Ptc and CiA for experiments have been obtained by polynomial fitting of the experimental profiles of at least 5 profiles for each condition. Next, we calculate the mean and the standard error for each parameter of the polynomial fitting. Finally, we construct a new polynomium using these average parameters, and represent the resulting curve as the average profile for each signal and each condition.

Experimental spatial profiles have been obtained by selecting a rectangular region in the sample. The width of the rectangular regions is 3 to 4 cell diameters, and the length is varied depending on the tissue being imaged. The spatial profile is obtained by average of the signal intensity across the width.

For the simulations, the individual profiles are obtained in a similar fashion as the experiments. In brief, the values of the expression levels for Hh, Ptc and CiA are averaged over the cells that are at a similar distance from the Hh source, transforming in this way information from a two-dimensional array of cells to a one-dimensional profile.

## ACKNOWLEDGEMENTS

Confocal microscopy was carried out at the ALMIA Platform (CABD). The Hh-GFP BAC strain is a gift of T. Kornberg (UCSF, USA) through I. Guerrero (CBM-SO, Madrid, EU).

## REFERENCES

Al-Saffar, Z.Y., Grainger, J.N.R., Aldrich, J., 1995. Effects of constant and fluctuating temperature 482 on development from egg to adult of drosophila melanogaster (meigen). Biology and Environment: Proceedings of the Royal Irish Academy 95B, 119–122.

Aza-Blanc, P., Ramirez-Weber, F.A., Laget, M.P., Schwartz, C., Kornberg, T.B., 1997. Proteolysis that is inhibited by hedgehog targets Cubitus interruptus protein to the nucleus and converts it to a repressor. Cell 89, 1043–1053.

Briscoe, J., Chen, Y., Jessell, T.M., Struhl, G., 2001. A hedgehog-insensitive form of patched provides evidence for direct long-range morphogen activity of sonic hedgehog in the neural tube. Mol Cell 7, 1279–1291.

Briscoe, J., Therond, P.P., 2013. The mechanisms of Hedgehog signalling and its roles in development and disease. Nat Rev Mol Cell Biol 14, 416–429.

Capdevila, J., Estrada, M.P., Sanchez-Herrero, E., Guerrero, I., 1994. The Drosophila segment polarity gene patched interacts with decapentaplegic in wing development. Embo J 13, 71–82.

Chen, W., Huang, H., Hatori, R., Kornberg, T.B., 2017. Essential basal cytonemes take up Hedgehog in the Drosophila wing imaginal disc. Development 144, 3134–3144.

Dietzl, G., Chen, D., Schnorrer, F., Su, K.C., Barinova, Y., Fellner, M., Gasser, B., Kinsey, K., Oppel, S., Scheiblauer, S., Couto, A., Marra, V., Keleman, K., Dickson, B.J., 2007. A genome-wide transgenic RNAi library for conditional gene inactivation in Drosophila. Nature 448, 151–156.

Dominguez, M., 1999. Dual role for Hedgehog in the regulation of the proneural gene atonal during ommatidia development. Development 126, 2345–2353.

Garcia-Morales, D., Navarro, T., Iannini, A., Pereira, P.S., Miguez, D.G., Casares, F., 2019. Dynamic Hh signalling can generate temporal information during tissue patterning. Development 146.

Haynie, J.L., Bryant, P.J., 1986. Development of the eye-antenna imaginal disc and morphogenesis of the adult head in Drosophila melanogaster. J Exp Zool 237, 293–308.

Heberlein, U., Wolff, T., Rubin, G.M., 1993. The TGF beta homolog dpp and the segment polarity gene hedgehog are required for propagation of a morphogenetic wave in the Drosophila retina. Cell 75, 913–926.

Ingham, P.W., Nakano, Y., Seger, C., 2011. Mechanisms and functions of Hedgehog signalling across the metazoa. Nat Rev Genet 12, 393–406.

Jarman, A.P., Sun, Y., Jan, L.Y., Jan, Y.N., 1995. Role of the proneural gene, atonal, in formation of Drosophila chordotonal organs and photoreceptors. Development 121, 2019–2030.

Kong, J.H., Siebold, C., Rohatgi, R., 2019. Biochemical mechanisms of vertebrate hedgehog signaling. Development 146.

Lee, R.T., Zhao, Z., Ingham, P.W., 2016. Hedgehog signalling. Development 143, 367–372.

Li, P., Markson, J.S., Wang, S., Chen, S., Vachharajani, V., Elowitz, M.B., 2018. Morphogen gradient reconstitution reveals Hedgehog pathway design principles. Science 360, 543–548.

Ma, C., Zhou, Y., Beachy, P.A., Moses, K., 1993. The segment polarity gene hedgehog is required for progression of the morphogenetic furrow in the developing Drosophila eye. Cell 75, 927–938.

Miguez, D.G., Garcia-Morales, D., Casares, F., 2020. Control of size, fate and time by the Hh morphogen in the eyes of flies. Curr Top Dev Biol 137, 307–332.

Miguez, D.G., Vanag, V.K., Epstein, I.R., 2007. Fronts and pulses in an enzymatic reaction catalyzed by glucose oxidase. Proc Natl Acad Sci U S A 104, 6992–6997.

Mohler, J., 1988. Requirements for hedgehog, a segmental polarity gene, in patterning larval and adult cuticle of Drosophila. Genetics 120, 1061–1072.

Mullor, J.L., Calleja, M., Capdevila, J., Guerrero, I., 1997. Hedgehog activity, independent of decapentaplegic, participates in wing disc patterning. Development 124, 1227–1237.

Nahmad, M., Stathopoulos, A., 2009. Dynamic interpretation of hedgehog signaling in the Drosophila wing disc. PLoS Biol 7, e1000202.

Petrova, R., Joyner, A.L., 2014. Roles for Hedgehog signaling in adult organ homeostasis and repair. Development 141, 3445–3457.

Roignant, J.Y., Treisman, J.E., 2009. Pattern formation in the Drosophila eye disc. Int J Dev Biol 53, 795–804.

Royet, J., Finkelstein, R., 1997. Establishing primordia in the Drosophila eye-antennal imaginal disc: the roles of decapentaplegic, wingless and hedgehog. Development 124, 4793–4800.

Sanicola, M., Sekelsky, J., Elson, S., Gelbart, W.M., 1995. Drawing a stripe in Drosophila imaginal disks: negative regulation of decapentaplegic and patched expression by engrailed. Genetics 139, 745–756.

Schindelin, J., Arganda-Carreras, I., Frise, E., Kaynig, V., Longair, M., Pietzsch, T., Preibisch, S., Rueden, C., Saalfeld, S., Schmid, B., Tinevez, J.Y., White, D.J., Hartenstein, V., Eliceiri, K., Tomancak, P., Cardona, A., 2012. Fiji: an open-source platform for biological-image analysis. Nat Methods 9, 676–682.

Schwartz, C., Locke, J., Nishida, C., Kornberg, T.B., 1995. Analysis of cubitus interruptus regulation in Drosophila embryos and imaginal disks. Development 121, 1625–1635.

Strigini, M., Cohen, S.M., 1997. A Hedgehog activity gradient contributes to AP axial patterning of the Drosophila wing. Development 124, 4697–4705.

Tabata, T., Kornberg, T.B., 1994. Hedgehog is a signaling protein with a key role in patterning Drosophila imaginal discs. Cell 76, 89–102.

Vollmer, J., Fried, P., Sanchez-Aragon, M., Lopes, C.S., Casares, F., Iber, D., 2016. A quantitative analysis of growth control in the Drosophila eye disc. Development 143, 1482–1490.

Wartlick, O., Julicher, F., Gonzalez-Gaitan, M., 2014. Growth control by a moving morphogen gradient during Drosophila eye development. Development 141, 1884–1893.

Wolpert, L., 1969. Positional information and the spatial pattern of cellular differentiation. J Theor Biol 25, 1–47.

